# ReTIF: Granularity-Aware Multitask Interaction Routing for RNA-Compound Interaction Prediction and Binding-Site Localization

**DOI:** 10.64898/2026.09.02.748859

**Authors:** Ao Chang, Heqin Zhu, Haobin Chen, Chenxi Wang, Xin Wang, Zeng Xiaoyu, Yilin Ding, Peng Xiong, S. Kevin Zhou

**Affiliations:** School of Biomedical Engineering, Division of Life Sciences and Medicine, University of Science and Technology of China (USTC), Hefei, Anhui, 230026, China; Medical Imaging, Robotics, Analytic Computing & Learning (MIRACLE) Lab, YRD-RIGHT, Suzhou Institute for Advance Research, USTC, Suzhou, Jiangsu, 215123, China; Biomedical Basic Research Center (BBRC) of Jiangsu, Suzhou, Jiangsu, 215123, China; Jiangsu Key Laboratory of Multimodal Digital Twin Technology, Suzhou, Jiangsu, 215123, China; State Key Laboratory of Precision and Intelligent Chemistry, Hefei, Anhui, 230026, China

## Abstract

RNA-targeted drug discovery requires both RNA–compound interaction prediction and nucleotide-level binding-site (BS) localization. These two tasks rely on different levels of interaction information: DTI prediction summarizes overall RNA-compound compatibility, whereas BS localization requires preserving nucleotide-level compound-associated signals. However, existing multitask models often use shared cross-modal representations for both tasks before task-specific prediction layers, which may limit their ability to preserve task-dependent interaction patterns. We therefore propose ReTIF(*Relation-enhanced Task-specific Interaction Framework*), which constructs separate cross-modal interaction representations for DTI prediction and BS localization before aggregation. ReTIF integrates multi-source RNA-compound representations from frozen RNA-FM, StructRFM, Mole-BERT, and MolFormer encoders, and builds separate interaction representations for DTI scoring and BS localization. The DTI branch captures global compatibility through aggregation, whereas the BS branch preserves nucleotide-compound resolution and enhances local evidence through relation-guided propagation with RNA structural and compound topological priors. An asymmetric DTI-derived compatibility signal provides global context to BS prediction while maintaining local evidence. Across five-fold evaluations under unseen pair, RNA, compound, and joint shifts with 13 base-lines, ReTIF has the highest mean in 13 of 16 scenario-metric combinations, including BS AUPR in all four settings.

## Introduction

RNA is increasingly recognized as a promising small-molecule therapeutic target, yet screening based on pair-level interaction scores provides limited guidance for downstream experimental validation (Connelly, Moon, and Schneekloth 2016; Warner, Hajdin, and Weeks 2018; Childs-Disney et al. 2022). RNA–compound recognition depends on local factors, including sequence motifs, secondary-structure contexts, loops, bulges, and compound-specific nucleotide environments. Therefore, effective computational models should not only estimate interaction likelihood but also identify nucleotide-level regions supporting molecular recognition (Warner, Hajdin, and Weeks 2018; Angelbello et al. 2018; Childs-Disney et al. 2022). Accordingly, pair-level drug–target interaction (DTI) prediction and nucleotide-level binding-site (BS) localization support complementary biological decisions: prioritizing promising RNA–compound pairs and identifying nucleotides for mechanistic investigation. Together, these predictions facilitate candidate screening, mutation analysis, structural interpretation, and compound optimization when experiments are costly.

Previous studies have explored pair-level interaction or affinity prediction, nucleotide-level binding-site localization, and joint prediction of both outputs (Huang et al. 2024; Chen et al. 2025; Zhu et al. 2025c; Bae and Nam 2026). Recent advances in RNA foundation models, structure-aware encoders, molecular graph representations, and chemical language models provide complementary sequence, structural, and chemical views. These developments make joint modeling attractive: DTI labels provide global compatibility signals, whereas BS labels provide fine-grained localization evidence. However, stronger unimodal representations do not directly determine how RNA and compound features should be combined for different prediction tasks.

We observe that DTI prediction and BS localization require different interaction representations. Although DTI and BS localization are biologically related, they require different evidence transformations: DTI prediction compresses distributed RNA–compound compatibility into a pair-level score, whereas BS localization preserves compound-conditioned evidence for each nucleotide (Caruana 1997). When both tasks share the same cross-modal interaction representation until prediction heads, global ranking and local localization are forced to optimize a common evidence space before task-specific aggregation. Such competing objectives can introduce representation interference, as demonstrated by previous multitask studies on optimization conflicts and adaptive sharing (Sener and Koltun 2018; Chen et al. 2018; Yu et al. 2020; Misra et al. 2016; Ma et al. 2018). This challenge becomes more pronounced under held-out RNA, compound, and joint shifts, where successful generalization requires transferring interaction evidence beyond observed entities. Therefore, we investigate whether task-specific interaction modeling should be introduced before or after cross-modal aggregation.

To address this challenge, we introduce ReTIF(*Relation-enhanced Task-specific Interaction Framework*), which moves task specialization from prediction heads to cross-modal interaction construction. ReTIF obtains complementary unimodal representations from frozen RNA-FM, StructRFM, Mole-BERT, and MolFormer encoders. Instead of sharing a unified interaction representation, ReTIF constructs independently parameterized interaction spaces for DTI prediction and BS localization before aggregation. The DTI branch aggregates global interaction evidence for pair-level scoring, whereas the BS branch preserves nucleotide– compound resolution and enhances local evidence through relation-guided propagation with structural and topological priors. An asymmetric DTI-derived compatibility signal provides global context to BS prediction without replacing nucleotide-resolved evidence. Overall, ReTIF shares unimodal representations while specializing cross-modal interaction modeling according to task granularity.

### Our main contributions are

- **A granularity-aware multitask interaction framework**. We identify interaction granularity mismatch as a key challenge in RNA–compound multitask prediction and construct task-specific cross-modal interaction spaces before aggregation.
- **A multi-source RNA–compound representation strategy**. We integrate frozen RNA-FM, StructRFM, Mole-BERT, and MolFormer representations to provide complementary sequence, structural, graph, and chemical views.
- **A relation-guided BS localization mechanism**. We enhance nucleotide-level prediction through intra-modal propagation guided by RNA structural and compound topological priors.

## Related Work

### RNA–Small-Molecule Interaction and Site Prediction

RNA–small-molecule modeling has developed along two related directions. Pair-level methods predict interaction or affinity: RSAPred combines RNA sequence features with ligand descriptors (Krishnan, Roy, and Gromiha 2024), DeepRSMA cross-fuses nucleotide- and atom-level features (Huang et al. 2024), and RNAsmol integrates molecular graphs, data augmentation, and attention fusion (Ma et al. 2025). These methods support pair-level screening through scores but lack compound-conditioned nucleotide labels.

Binding-site methods instead localize interacting nucleotides. RLBind combines global RNA information with local sequence and structure features, whereas RLsite integrates pretrained RNA language models with graph attention (Wang et al. 2023; Sun et al. 2025); PCN-RNAsite combines position-specific and complex-network features (Zhang, Xiao, and Kong 2024); and MVRBind and RNABind incorporate multi-view structural or geometric representations (Chen et al. 2025; Zhu et al. 2025c). However, they do not jointly optimize pair-level DTI and compound-conditioned BS prediction. SMRTnet links the tasks through gradient-based response maps without an explicit BS head (Fei et al. 2026), whereas GerNA-Bind uses multistate RNA–ligand geometry to model binding specificity and structure-level sites (Xia et al. 2025). DeepRNA-DTI is the closest task-matched prior: it jointly supervises DTI and BS prediction and uses DTI output to guide BS localization (Bae and Nam 2026). ReTIFretains this asymmetric DTI-to-BS link but moves task specialization into interaction construction, building pooled DTI and nucleotide– compound-resolved BS tensors before aggregation.

### Architectures and Foundation Representations

Protein DTI architectures provide references for compound– target prediction. DeepDTA encodes drug SMILES and target sequences with convolutional networks (Öztürk, Özgür, and Ozkirimli 2018); GraphDTA introduces molecular graph representations (Nguyen et al. 2021); and GraphATT-DTA adds attention-based target–compound interaction modeling (Bae and Nam 2023). Direct adaptation to RNA, however, does not address RNA-specific structure, nucleotide-level localization, or preservation of nucleotide–compound resolution under distribution shifts.

Pretrained encoders provide complementary priors: RNA-FM models RNA sequence context (Chen et al. 2022), StructRFM adds structure-guided RNA pretraining (Zhu et al. 2025a), Mole-BERT captures graph-local molecular topology (Xia et al. 2023), and MolFormer learns SMILES representations (Ross et al. 2022). BPfold, NCfold, and IRE-Seek further combine sequence representations with physical or learned structural priors for secondary-structure, non-canonical base-pair, and IRES prediction, respectively (Zhu et al. 2025b, 2026; Zhang et al. 2025). These advances improve unimodal representations but do not determine when RNA–compound evidence should be specialized or pooled, motivating task-specific interaction construction.

### Task-Aware Interaction Routing

Site-aware RNA–compound prediction couples DTI and BS labels at different resolutions. Multitask learning provides an inductive bias (Caruana 1997), while cross-stitch and multigate architectures learn task-specific sharing (Misra et al. 2016; Ma et al. 2018). Competing objectives can nevertheless strain shared representations (Sener and Koltun 2018; Chen et al. 2018; Yu et al. 2020), motivating separate evidence construction for global and local outputs. Unlike late branching at prediction heads, ReTIF specializes the cross-modal interaction stage: shared tokens feed separate operators for globally pooled DTI evidence and nucleotide-resolved BS evidence. The BS pathway retains an asymmetric forward dependency through a DTI-head-derived compatibility gate computed within its own forward pass.

### Problem Formulation

Given an RNA sequence 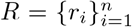 and a compound *C*, we consider two coupled prediction tasks: RNA–compound interaction prediction and nucleotide-level binding-site (BS) localization. The drug–target interaction (DTI) task determines whether the RNA and compound interact, producing a pair-level probability *ŷ*_DTI_ ∈ [0, 1]. The BS task identifies compound-associated nucleotides and outputs residue-level probabilities **ŷ**_BS_ ∈ [0, 1]^*n*^. Although sharing the same RNA–compound pair, the two tasks require different cross-modal evidence granularities: DTI relies on aggregated molecular compatibility, whereas BS localization preserves nucleotide–compound interactions for residue-level prediction. Therefore, we formulate RNA–compound prediction as a coupled multi-task learning problem with task-specific interaction modeling. Training uses pair-level DTI labels and nucleotide-level BS annotations. BS supervision is applied only to interacting pairs, with padding positions masked during BS loss computation and evaluation. Following previous benchmarks, we evaluate both tasks under four held-out generalization settings: unseen RNA–compound pairs, unseen RNAs, unseen compounds, and jointly unseen RNAs and compounds. DTI performance is measured using AUC and AUPR for pair-level ranking, while BS AUC/AUPR are computed within each valid interacting pair over non-padding nucleotides and averaged across pairs.

### Overview

#### Methodology

RNA–compound interaction prediction and binding-site localization require different cross-modal granularities. DTI prediction benefits from aggregated molecular compatibility, whereas BS localization requires preserving nucleotide– compound interaction resolution before prediction.

As illustrated in Figure 1, ReTIF(*Relation-enhanced Task-specific Interaction Framework*) addresses this granularity mismatch by constructing task-specific interaction fields before cross-modal pooling. Given RNA and compound inputs, ReTIFfirst builds multi-source token representations using frozen RNA-FM, StructRFM, Mole-BERT, and MolFormer encoders. Instead of forcing both tasks to share a unified interaction space, it constructs independent DTI and BS interaction tensors with different aggregation strategies. The DTI pathway performs global compatibility aggregation for pair prediction, while the BS pathway preserves nucleotide-level resolution and enhances local evidence through relation-guided propagation. An asymmetric DTI-to-BS transfer provides compatibility context for BS localization, while molecular priors weakly regularize intra-molecular propagation.

**Figure 1:**
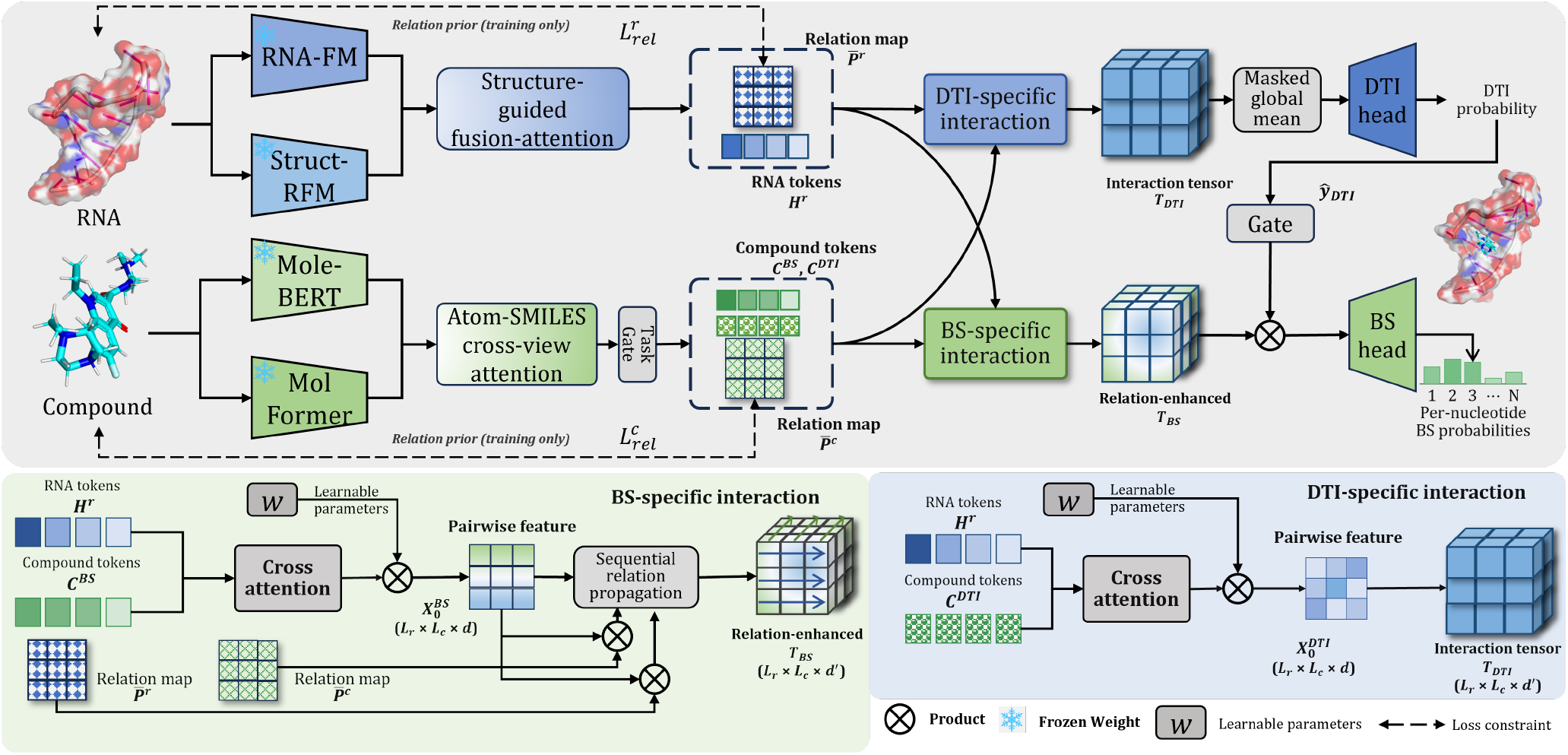
Overview of ReTIF. Shared RNA and compound token features feed task-specific DTI and BS interaction tensors. The DTI branch produces a global pair score, whereas the BS branch retains nucleotide–compound resolution for relation-guided localization. The gate denotes an asymmetric DTI-derived compatibility signal computed within the BS forward pass.

### Multi-Source Representation Encoding

#### RNA representation

We extract RNA representations from RNA-FM and StructRFM, which capture sequence-level and structure-aware information, respectively. Let *L*_*r*_ and *L*_*c*_ denote the numbers of RNA and compound tokens, respectively, and let *d* be the shared hidden dimension. After projection into a shared latent space, RNA-FM tokens query StructRFM tokens through cross-attention. The attended structural update is added to the RNA-FM representation and layer-normalized to produce **H**^*r*^. This fusion preserves one representation for each nucleotide while incorporating structural context before RNA–compound interaction construction. The corresponding attention weights are used as intra-RNA propagation signals in later BS refinement.

#### Compound representation

For compounds, Mole-BERT provides atom-level graph tokens 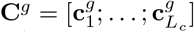, while MolFormer provides SMILES-level representations. To integrate graph topology and sequential chemical semantics, each atom token attends to the SMILES tokens. The attended update is added to its graph representation and layer-normalized, yielding 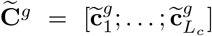. The resulting atom-level representations preserve both molecular topology and SMILES-derived contextual information for subsequent task-specific interaction modeling.

### Task-conditioned Modality Fusion

Different prediction objectives may rely on different molecular views. DTI prediction benefits from global molecular compatibility, whereas BS localization requires fine-grained chemical context around binding nucleotides.

Therefore, ReTIF learns task-specific compound representations by dynamically weighting graph and SMILES views:

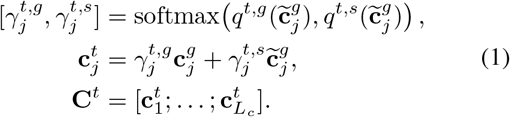

Here, *t* dti, bs and *q*^*t*,·^ denotes a scalar projection.

The fused compound tokens are then used by independent DTI and BS interaction modules.

### Task-Specific Interaction Construction

Existing multitask RNA–compound models typically share a unified cross-modal representation until prediction heads. However, DTI prediction and BS localization require different evidence granularities: DTI relies on globally aggregated molecular compatibility, whereas BS prediction requires preserving nucleotide–compound interaction resolution.

To address this mismatch, ReTIF constructs task-specific interaction fields before cross-modal pooling. Given the shared RNA representation **H**^*r*^ and task-specific compound representations **C**^*t*^, where *t* ∈ {dti, bs}, each task independently builds its own cross-modal evidence space.

For each task, an independent cross-attention module lets **H**^*r*^ query **C**^*t*^. Its compound-aware update is passed through dropout, added residually to **H**^*r*^, and layer-normalized to produce task-conditioned RNA tokens **H**^*r,t*^; we denote its *i*-th token by 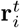. These tokens preserve different conditioned contexts for DTI and BS prediction. We then construct an RNA–compound interaction field through bilinear token interactions. For an RNA token *i* and a compound token *j*, the interaction affinity and feature are defined as:

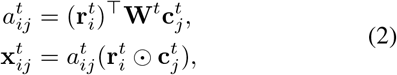

where **W**^*t*^ denotes a task-specific bilinear projection and ⊙ represents element-wise multiplication.

Stacking these pairwise features yields 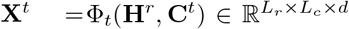, where Φ_*t*_ denotes the complete task-specific interaction construction above. Instead of sharing this tensor across tasks, ReTIF maintains independent DTI and BS interaction fields:

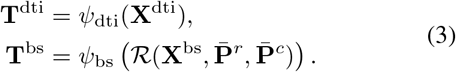

Here, *ψ*_dti_ and *ψ*_bs_ are independent task-specific feature projections, and ℛ denotes the sequential relation propagation defined below.

The DTI field is later aggregated for pair-level prediction, whereas the relation-refined BS field preserves nucleotide– compound resolution for localization.

### Relation-guided BS Evidence Propagation

The raw BS interaction field **X**^bs^ preserves nucleotide– compound resolution, but each pairwise feature initially captures only local compatibility. ReTIF therefore refines this field with intra-molecular context derived from nucleotide dependencies and molecular topology; this relation propagation is applied only to the BS pathway.

We derive propagation operators from the attention interactions learned during representation encoding. Let **P**^*r*^ denote the structure-guided RNA attention matrix and **A**^*s*^ the head-averaged atom–SMILES attention matrix. Using row normalization RN(·) over valid positions, the RNA and compound propagation operators are

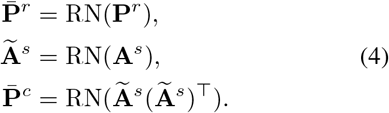

These operators propagate learned intra-molecular context along the RNA and compound axes rather than serving as post-hoc explanations. Interaction labels remain responsible for determining cross-modal compatibility.

Given propagation strengths *α*_*r*_ and *α*_*c*_, the BS interaction tensor is refined sequentially along RNA and compound dimensions:

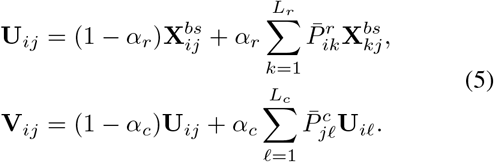

Together, these updates define 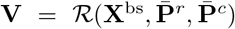. The refined tensor incorporates neighboring nucleotide and molecular context while preserving nucleotide–compound resolution for subsequent BS prediction.

### Asymmetric DTI-to-BS Transfer

DTI prediction provides global RNA–compound compatibility, which can offer useful context for BS localization.

Therefore, ReTIFintroduces a one-way DTI-to-BS compatibility transfer.

The DTI branch applies masked mean pooling, denoted by Pool, over the valid positions of **T**^dti^. It then uses the classifier *h*_dti_ followed by a sigmoid function to obtain the interaction probability *ŷ*_DTI_.

During BS prediction, the DTI interaction operator Φ_dti_ is reused with the BS-conditioned compound representation. Its compatibility score gates the relation-refined BS tensor:

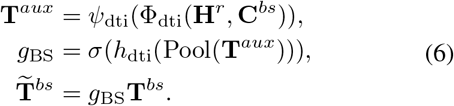

For each nucleotide, the BS head *h*_bs_ scores its compound-conditioned interaction features, takes their masked mean over valid compound positions, and applies a sigmoid function to obtain the binding-site probability *ŷ*_BS,*i*_.

This asymmetric transfer provides global compatibility context for BS refinement while preserving nucleotide– compound resolution; the forward design contains no reverse BS-to-DTI modulation.

### Weak Molecular Prior Regularization

To incorporate molecular structural knowledge into BS refinement, ReTIFapplies weak prior regularization to the learned propagation operators.

For RNA, the prior matrix combines ViennaRNA-predicted secondary-structure adjacency with local nucleotide proximity. For compounds, the prior matrix is derived from molecular graph connectivity. These priors constrain intra-molecular information propagation without assuming explicit RNA–compound contact annotations.

The regularization loss is defined as:

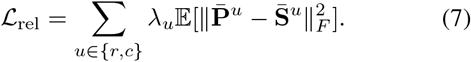

Here 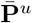 denotes the learned propagation operator, 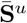 the corresponding molecular prior, and *λ*_*u*_ its relation-specific weight. The expectation averages the penalty over valid prior rows and minibatch examples. The weak constraint guides intra-molecular refinement while allowing interaction labels to determine cross-modal compatibility. The overall objective is

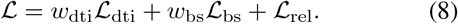

Detailed loss definitions and weights are provided in the supplementary material.

### Experiments

#### Experimental Setup

##### Dataset and protocol

We evaluate on the DeepRNA-DTI benchmark (Bae and Nam 2026), comprising 586 RNAs and 2,928 compounds with pair-level DTI and nucleotide-level BS labels. We follow the official five-fold protocol under *unseen pair, unseen RNA, unseen compound*, and *unseen both* splits. All test metrics are averaged across folds.

##### Baselines and metrics

We compare ReTIFwith 13 interaction, BS-localization, adapted protein-DTI, and classical machine-learning baselines. DTI AUC/AUPR are computed over held-out RNA–compound pairs. BS AUC/AUPR are computed over nucleotides within each test pair containing annotated sites and macro-averaged across eligible pairs. Single-task methods are evaluated only on outputs. We rerun all baselines under the same five-fold protocol.

##### Training

RNA-FM, StructRFM, Mole-BERT, and MolFormer remain frozen. ReTIF is trained with AdamW (Loshchilov and Hutter 2019) for 100 epochs, with non-interacting pairs and padded nucleotides excluded from the BS loss. Baseline adaptations and hyperparameter settings are provided in the supplementary material.

#### Main Results

Table 1 compares both output granularities across four held-out scenarios. ReTIF has the highest five-fold mean in 13 of the 16 scenario–metric combinations: six of eight DTI results and seven of eight BS results. The clearest pattern in the means occurs in nucleotide-level localization. This pattern does not extend uniformly to every metric. ReTIF has the highest mean BS AUPR in all four scenarios and the highest mean BS AUC in three. The ReTIF–DeepRNA-DTI mean differences in BS AUC/AUPR are 0.015/0.020, 0.039/0.027, 0.008/0.013, and 0.068/0.040 under *unseen pair, unseen RNA, unseen compound*, and *unseen both*, respectively. Relative to the highest BS-only AUPR mean in each scenario, the corresponding mean margins are 0.117, 0.044, 0.179, and 0.098. These AUPR differences are especially informative because binding nucleotides are sparse and AUPR emphasizes ranking quality for the positive class. The only BS metric not led by ReTIF is *unseen RNA* AUC, where RL-site scores 0.614 versus 0.581; nevertheless, ReTIF has the highest mean AUPR and higher mean BS AUC/AUPR than DeepRNA-DTI in this split.

**Table 1:** Main results across four different scenarios. DTI AUC, DTI AUPR, BS AUC, and BS AUPR are 5-fold means. Best and <u>second-best</u> in each column. “–” = not applicable.

| Model | Type | unseen pair |  |  |  | unseen RNA |  |  |  | unseen compound |  |  |  | unseen both |  |  |  |
| --- | --- | --- | --- | --- | --- | --- | --- | --- | --- | --- | --- | --- | --- | --- | --- | --- | --- |
|  |  | DTI |  | BS |  | DTI |  | BS |  | DTI |  | BS |  | DTI |  | BS |  |
|  |  | AUC | AUPR | AUC | AUPR | AUC | AUPR | AUC | AUPR | AUC | AUPR | AUC | AUPR | AUC | AUPR | AUC | AUPR |
| Logistic Reg. | Classical | 0.557 | 0.551 | – | – | 0.534 | 0.533 | – | – | 0.499 | 0.480 | – | – | 0.510 | 0.496 | – | – |
| SVM (Linear) | Classical | 0.537 | 0.515 | – | – | 0.515 | 0.508 | – | – | 0.500 | 0.476 | – | – | 0.487 | 0.499 | – | – |
| DeepDTA | Protein DTI | 0.585 | 0.605 | – | – | 0.539 | <u>0.571</u> | – | – | 0.538 | 0.511 | – | – | 0.469 | 0.493 | – | – |
| GraphDTA | Protein DTI | 0.720 | 0.714 | – | – | <u>0.553</u> | 0.550 | – | – | 0.585 | 0.555 | – | – | 0.511 | 0.538 | – | – |
| GraphATT-DTA | Protein DTI | 0.721 | 0.704 | – | – | <u>0.553</u> | 0.550 | – | – | <u>0.610</u> | 0.564 | – | – | <u>0.526</u> | 0.528 | – | – |
| RSAPred | RNA interact | 0.631 | 0.598 | – | – | 0.518 | 0.514 | – | – | 0.517 | 0.474 | – | – | 0.463 | 0.487 | – | – |
| DeepRSMA | RNA interact | 0.514 | 0.539 | – | – | 0.509 | 0.501 | – | – | 0.484 | 0.469 | – | – | 0.523 | <u>0.539</u> | – | – |
| RNABind | BS-only | – | – | 0.724 | 0.530 | – | – | 0.552 | 0.397 | – | – | 0.649 | 0.511 | – | – | 0.523 | 0.608 |
| RLBind | BS-only | – | – | <u>0.833</u> | 0.660 | – | – | <u>0.594</u> | 0.448 | – | – | 0.755 | 0.663 | – | – | 0.509 | 0.594 |
| RLsite | BS-only | – | – | 0.831 | 0.656 | – | – | **0.614** | 0.463 | – | – | 0.716 | 0.645 | – | – | <u>0.666</u> | 0.691 |
| MVRBind | BS-only | – | – | 0.733 | 0.541 | – | – | 0.552 | 0.400 | – | – | 0.673 | 0.539 | – | – | <u>0.539</u> | 0.591 |
| PCN-RNAsite | BS-only | – | – | 0.736 | 0.523 | – | – | 0.580 | 0.394 | – | – | 0.638 | 0.480 | – | – | 0.563 | 0.628 |
| DeepRNA-DTI | Multitask | <u>0.729</u> | <u>0.725</u> | 0.828 | <u>0.757</u> | 0.552 | 0.554 | 0.542 | <u>0.480</u> | **0.618** | **0.579** | 0.849 | 0.829 | 0.514 | 0.495 | 0.627 | <u>0.749</u> |
| ReTIF | Multitask | **0.732** | **0.731** | **0.843** | **0.777** | **0.575** | **0.586** | 0.581 | **0.507** | 0.602 | <u>0.574</u> | **0.857** | **0.842** | **0.576** | **0.580** | **0.695** | **0.789** |

Pair-level DTI results depend more strongly on the type of distribution shift. The ReTIF–DeepRNA-DTI mean differences in DTI AUC/AUPR are 0.003/0.006 for *unseen pair*, 0.023/0.032 for *unseen RNA*, and 0.062/0.085 for *unseen both*. Under joint shift, its mean DTI AUC and AUPR are also higher than the strongest competing values by 0.050 and 0.041, respectively. The trend reverses for *unseen compound*: DeepRNA-DTI reaches 0.618/0.579 AUC/AUPR versus 0.602/0.574 for ReTIF. The BS mean differences are positive in all four shifts; the DTI mean differences are largest under RNA and joint shifts and negative under compound shift.

#### Ablation Study

We next separate the effects associated with the multi-source backbone, the cross-task sharing boundary, and the prior-guided relation-map package. Figure 2(a–d) compares task-specific interaction routing, the relation-map package, and weak map supervision; complete five-fold routing, relation-map, and backbone results are provided in the supplementary material. G2 replaces task-specific tensors with a shared interaction stage and removes the BS relation-map package. G1 retains task-specific routing but removes the same relation maps. Because both variants exclude propagation, their contrast narrows the architectural difference to shared versus task-specific interaction construction. G1 has higher mean BS AUC/AUPR than G2 by 0.046/0.059 for *unseen pair* and 0.065/0.043 for *unseen both*, while differences are negligible or mixed under the two single-entity shifts. The Full–G2 mean DTI AUC/AUPR differences are positive in three of four scenarios and reach 0.062/0.061 under *unseen both*.

**Figure 2:**
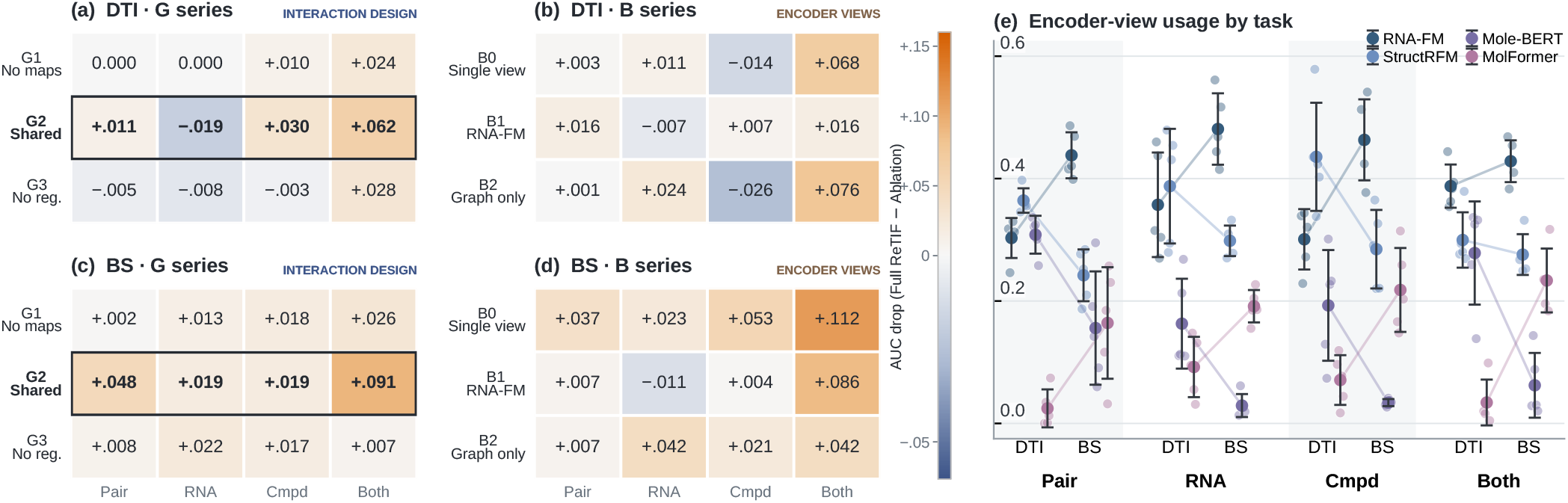
Ablations and view diagnostics across matched folds. (a)–(d) report mean AUC drops relative to Full, where positive values favor Full and outlines denote G2. G1 removes relation maps, G2 shares the interaction tensor, and G3 removes map regularization. B0 retains only RNA-FM and Mole-BERT, B1 removes StructRFM, and B2 removes MolFormer; all backbone variants omit relation maps. (e) shows normalized post-hoc encoder-view usage on each task set rather than learned gates.

Restoring the relation-map package to G1 produces positive mean BS differences in all scenarios, with Full–G1 gaps ranging from 0.001 to 0.027 across BS AUC and AUPR. The Full–G3 mean BS gaps range from 0.006 to 0.022 and are smaller than the routing-related gaps under pair and joint shifts. To examine the multi-source backbone without relation maps, we compare G1 with B0–B2, which also omit the relation-map package. B0 retains only RNA-FM and Mole-BERT; the G1–B0 mean BS AUC/AUPR differences are 0.035/0.044, 0.009/0.013, 0.035/0.055, and 0.085/0.063 under *unseen pair, unseen RNA, unseen compound*, and *unseen both*, respectively. DTI mean differences are mixed, with B0 higher under compound shift. The added views contribute asymmetrically: removing StructRFM lowers BS metrics under pair and joint shifts but raises them under single-entity shifts, whereas removing MolFormer lowers BS AUC in all scenarios and BS AUPR in three while improving compound-shift DTI. The largest G1–G2 mean gaps occur under pair and joint shifts; the relation-map gaps are smaller and positive across BS metrics, while the structural and SMILES views show shift-dependent differences.

#### Analysis

##### Task-specific encoder-view usage

Figure 2(e) summarizes task-specific use of the four encoder views through post-hoc forward-activation diagnostics. These normalized values are not four learned gates: RNA shares reflect relative feature norms, whereas compound shares average the learned graph/SMILES gate weights in Eq. (1) over valid atoms. On the BS evaluation set, RNA-FM is larger in all four splits (0.428–0.481), while StructRFM is larger on the DTI evaluation set in three. Compound diagnostics are graph-dominant for DTI, but string-dominant for BS under RNA, compound, and joint shifts. Fold-level shares and normalization details for all five folds appear in the supplementary material.

##### Spatial organization of case-level predictions

Figure 3 provides a qualitative comparison of BS localization across four held-out settings: unseen pair (8I45), unseen RNA (7REX), unseen compound (2EEU), and unseen both (1TOB). It examines whether predicted sites recover ligand-associated nucleotide organizations from sequence and three-dimensional views. LigPlot summarizes local contact neighborhoods, while PyMOL shows the complete RNA surface and the spatial organization of predicted sites. Ground-truth BS labels follow the 10 Å criterion, and predictions are thresholded at 0.5. The sequence plots further show prediction continuity and coverage along the RNA sequence.

**Figure 3:**
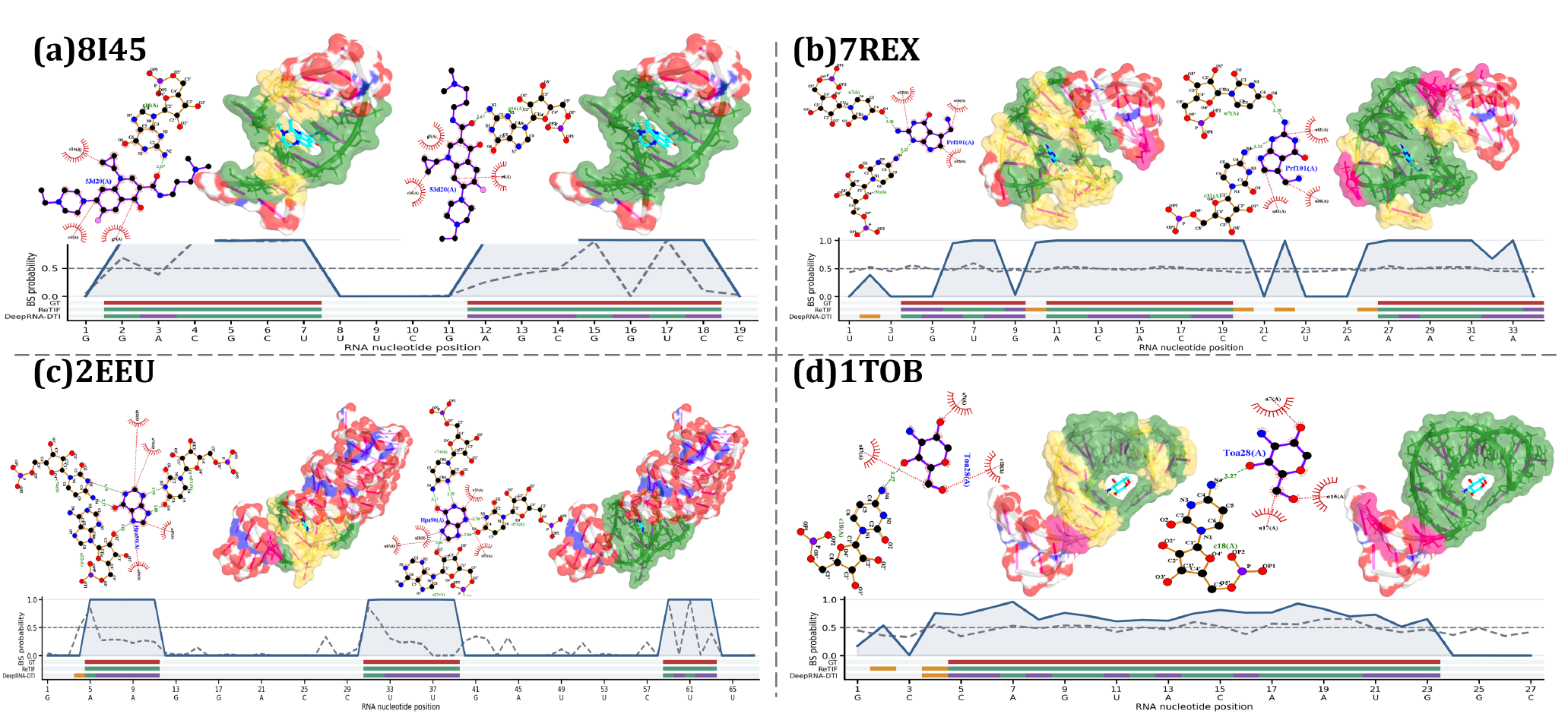
BS predictions for mapped PDB/ligand-component pairs: (a) *unseen pair*, 8I45/53D; (b) *unseen RNA*, 7REX/PRF; (c) *unseen compound*, 2EEU/HPA; and (d) *unseen both*, 1TOB/TOA. LigPlot shows contacts; PyMOL compares DeepRNA-DTI (left) and ReTIF(right) over the complete RNA surface. Ground-truth BS labels use the 10 Å criterion and predictions use a 0.5 threshold; TP, FP, and FN are green, pink, and yellow, with the ligand in cyan. Sequence plots show ReTIF(solid blue) and DeepRNA-DTI (dashed gray); strips show ground truth and outcomes.

In 8I45 (Fig. 3(a)) and 2EEU (Fig. 3(c)), labeled nucleotides span multiple separated sequence intervals that converge around KG022 (53D) and hypoxanthine (HPA), respectively, in three-dimensional space. ReTIF better recovers these discontinuous regions and produces predictions that better overlap with ligand-associated regions, whereas DeepRNA-DTI generates fragmented predictions with missing positions around ligand-associated areas. These cases suggest that ReTIF better captures BS evidence distributed across separated sequence regions.

In 7REX (Fig. 3(b)), ReTIF captures most labeled regions associated with preQ1 (PRF), including the 11–19 interval, while retaining limited FP positions. DeepRNA-DTI produces sparser predictions and misses more annotated sites. In 1TOB (Fig. 3(d)), ReTIF maintains predictions above the decision threshold across the continuous 5–23 region associated with the selected tobramycin sugar component (TOA) and largely avoids FN positions within the annotated interval, whereas DeepRNA-DTI exhibits alternating TP and FN predictions.

Overall, ReTIF improves the spatial coherence, coverage, and continuity of BS predictions rather than simply increasing correctly classified nucleotides. This behavior is consistent with relation-enhanced BS evidence propagation, which models nucleotide-level dependencies along RNA and compound axes. Since BS labels are defined by a 10 Å distance cutoff, these visualizations assess the spatial organization of classification outcomes rather than directly validating molecular contacts or binding mechanisms.

## Conclusion

ReTIF shares frozen multi-source RNA and compound representations while constructing independently parameterized DTI and BS interaction tensors before pooling. The DTI pathway captures global pairwise compatibility, whereas the BS pathway preserves nucleotide–compound resolution and refines local evidence through relation maps guided by structural and topological priors. An asymmetric DTI-derived gate provides global compatibility context to the BS pathway without replacing its local interaction modeling. Across four held-out scenarios and 13 baselines, ReTIF achieves the best five-fold mean in 13 of 16 scenario–metric entries, including BS AUPR across all scenarios.

